# *Brucella ovis* cysteine biosynthesis contributes to peroxide stress survival and fitness in the intracellular niche

**DOI:** 10.1101/2020.12.22.424080

**Authors:** Lydia M. Varesio, Aretha Fiebig, Sean Crosson

## Abstract

*Brucella ovis* is an ovine intracellular pathogen with tropism for the male genital tract. To establish and maintain infection, *B. ovis* must survive stressful conditions inside host cells, including low pH, nutrient limitation, and reactive oxygen species. These same conditions are often encountered in axenic cultures during stationary phase. Studies of stationary phase may thus inform understanding of *Brucella* infection biology, yet the genes and pathways that are important in *Brucella* stationary phase physiology remain poorly defined. We measured fitness of a barcoded pool of *B. ovis* Tn-*himar* mutants as a function of growth phase and identified *cysE* as a determinant of fitness in stationary phase. CysE catalyzes the first step in cysteine biosynthesis from serine, and we provide genetic evidence that two related enzymes, CysK1 and CysK2, function redundantly to catalyze cysteine synthesis at steps downstream of CysE. Deleting either *cysE* (Δ*cysE*) or both *cysK1* and *cysK2* (Δ*cysK1* Δ*cysK2*) results in premature entry into stationary phase, reduced culture yield and sensitivity to exogenous hydrogen peroxide. These phenotypes can be chemically complemented by cysteine or glutathione. Δ*cysE* and Δ*cysK1* Δ*cysK2* strains have no defect in host cell entry *in vitro* but have significantly diminished intracellular fitness between 2 and 24 hours post infection. Our study has uncovered unexpected redundancy at the CysK step of cysteine biosynthesis in *B. ovis*, and demonstrates that cysteine anabolism is a determinant of peroxide stress survival and fitness in the intracellular niche.

## Introduction

*Brucella* spp. are intracellular pathogens that have numerous mechanisms to contend with host-generated stressors and exploit host resources for growth. Within the host, brucellae are subject to nutrient limitation (1), phagosomal acidification (2), and direct attack from reactive oxygen and reactive nitrogen species (3) originating from the host-derived respiratory burst (4, 5). Dozens of genes involved in oxidative stress responses, acid stress responses, nutrient assimilation, and respiration have been implicated in the biology of *Brucella* infection (6). More recent studies have defined a role for the general stress response pathway in mitigation of multiple chemical stressors *in vitro* and in maintenance of chronic infection *in vivo* (7, 8). However, relatively little is known about the mechanisms *Brucella* spp. use to adapt to stresses encountered in axenic cultures during stationary phase. The study of stationary phase has the potential to inform the discovery of genes that influence infection, intracellular replication, and survival (9) as there are postulated parallels between stationary phase physiology and the physiologic state of *Brucella* in the intracellular niche (1).

We sought to develop an approach to identify genes involved in stationary phase physiology in the ovine pathogen, *Brucella ovis. B. ovis* (10) is an understudied member of the *Brucella* clade that has a number of distinguishing genomic features (11). It is one of two naturally rough species among the classical *Brucella* group (12) and is the only species of this group that is non-zoonotic. The host environment inhabited by *B. ovis* is quite restricted: it is typically sexually transmitted and has a specific tropism for the male genital tract in rams (13–15). We previously constructed a randomly barcoded (RB) library of *B. ovis* Tn-*himar* mutants (16) and in this present study we set out to develop this barcoded mutant library as a tool to identify *B. ovis* genes with fitness defects in stationary phase culture. We measured the relative fitness of RB Tn-*himar* mutants as a function of growth phase in a complex medium and discovered that disruption of the cysteine biosynthesis gene, *cysE*, resulted in the largest stationary phase fitness defect in our experiment. Thus, our screen provided evidence for a link between a sulfur assimilation/cysteine biosynthesis pathway and stationary phase physiology.

Recent studies have significantly advanced our understanding of the roles of carbon and nitrogen metabolism in *Brucella* physiology and infection (17, 18), but our knowledge of sulfur metabolism in *Brucella* spp. is comparatively limited. This study defines the role of a sulfur metabolism pathway in the growth, stress survival, and infection biology of *Brucella*. Specifically, we show that *cysE*-dependent biosynthesis of cysteine influences culture yield and stationary phase entry in *B. ovis*. We further provide genetic evidence that two related CysK-family enzymes, CysK1 and CysK2, function redundantly at a step downstream of CysE to produce cysteine. *B. ovis* strains lacking *cysE* or both *cysK1* and *cysK2* are compromised in peroxide stress survival and in colonization of mammalian host cells.

## Results

### *B. ovis cysE* Tn-*himar* mutant strains have a fitness defect in stationary phase

We inoculated ≈ 1.5 × 10^9^ *B. ovis* RB Tn-*himar* strains into Brucella Broth in triplicate and collected samples at intervals throughout the growth curve: 0.05, 0.12, 0.9 and 2.4 OD_600_ (corresponding to early logarithmic, logarithmic, late logarithmic, and stationary phase). Barcodes were PCR amplified, sequenced, and tallied as previously described (19) to assess the relative abundance of each mutant strain in each sample. Our analysis yielded composite fitness scores for 2638 of 3391 annotated genes in *B. ovis* **(Data Set 1)**. Data for 118 mutants that exceeded a t-like test significance threshold ≥ 4 are presented in **Fig. S1 (**see **Materials and Methods)**. We observed the largest relative fitness score changes at OD_600_ = 2.4 (i.e., stationary phase) in this dataset.

To more rigorously assess mutants with fitness values that varied as a function of growth phase, we further filtered the genes to include only those with a fitness score ≥ |1| in at least one timepoint (see **Materials and Methods**). We hierarchically clustered the 64 genes that passed this cutoff (**Fig. 1A, Data Set 1**) and divided these clustered genes into four groups that displayed different fitness patterns throughout the growth curve (**Fig. 1B**). Mutations in group 1 genes resulted in no fitness defect during exponential growth, but a fitness defect at OD_600_ = 2.4 (i.e. stationary phase). Genes in group 2 had negative fitness scores throughout the growth curve. These two groups contained the majority of mutants. Group 3 (four genes) had positive fitness scores in log phase and a negative fitness score in stationary phase, while group 4 (three genes) had null or positive fitness scores at all phases of the growth curve. We clustered these genes by predicted functional category (**Fig. S2A, Data Set 2**) and found that genes encoding purine metabolism enzymes and tRNA modification enzymes were enriched in group 1. However, the gene with the lowest fitness score in stationary phase, *BOV_RS06060* (old locus tag *BOV_1224*), is annotated as serine-O-acetyltransferase (*cysE*) (**Fig. 1** and **Fig. S2B**). As such, we chose to further characterize the function of *cysE* in *B. ovis*.

**Figure 1.**
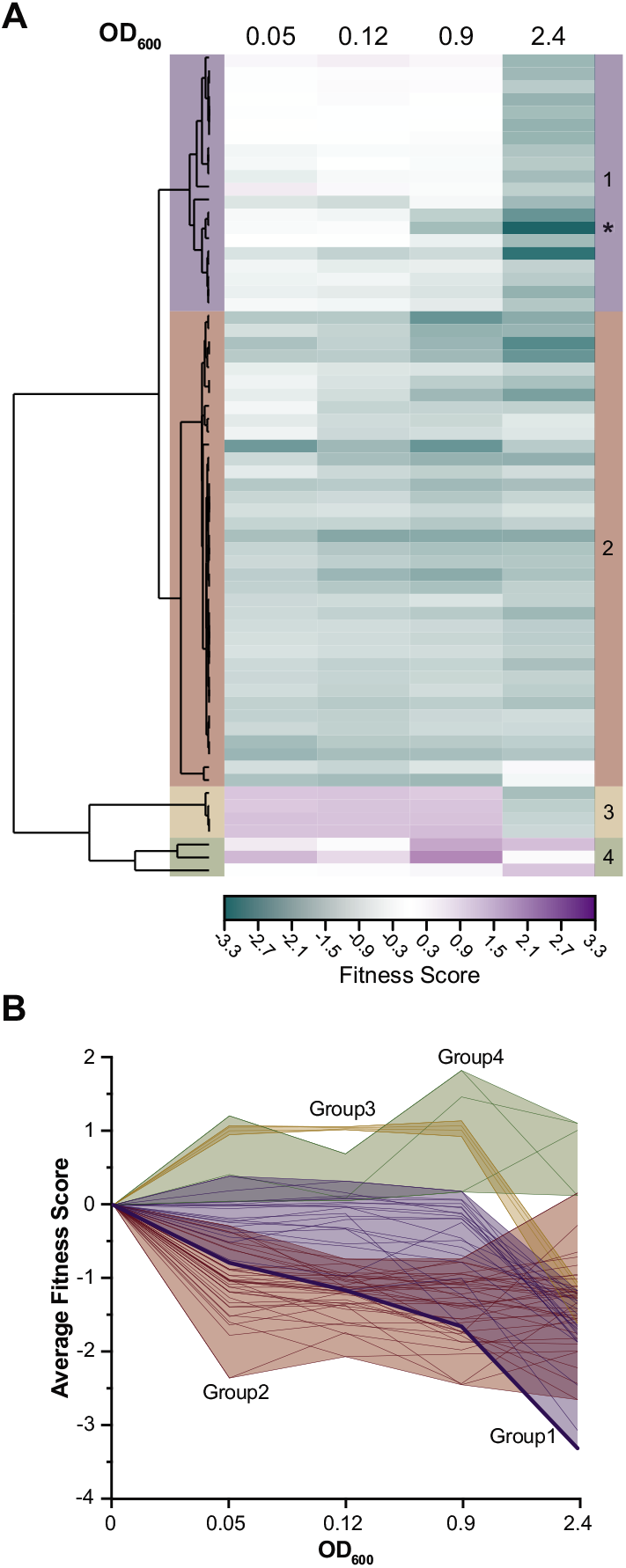
Assessment of *B. ovis* mutant strain fitness as a function of growth phase identifies *cysE* as a determinant of stationary phase fitness. **A)** Heat map showing mean fitness scores (n = 3) of *B. ovis* mutant strains harboring barcoded transposon insertions in non-essential genes. Each row represents a gene, and each column is a point during the growth curve in BB (OD_600_). Genes with a t-like significance score ≥ |4| and fitness value ≥ |1| in at least one timepoint are included in the heat map. Genes were hierarchically clustered (left), which yielded four main groups (right). The fitness profile of strains harboring RB Tn-*himar* insertions in *cysE* is marked with an asterisk on the right of the heat map. **B)** Alternative representation of data shown in **A**, where the average fitness score for each mutant (gene) is plotted as a function of timepoint. Individual lines represent genes, shaded area delimits the max and minimum values within that group. *cysE* is presented as a thick blue line. Group 1, blue; Group 2, red; Group 3, yellow; Group 4, green.

### *B. ovis* Δ*cysE* enters stationary phase prematurely and has reduced culture yield *in vitro*

*B. ovis* CysE has high sequence identity (52%) and similarity (73%) with the well-characterized CysE enzymes of *Escherichia coli* and *Salmonella enterica* (20), and is clearly classified as CysE in the NCBI conserved domain database (E-value = 4.3e^-122^, https://www.ncbi.nlm.nih.gov/Structure/cdd). This protein is therefore predicted to execute the initial step in cysteine biosynthesis, specifically the addition of an acetyl group from acetyl-CoA to serine, producing O-acetylserine (**Fig. 2A**). To confirm the *cysE* stationary phase phenotype observed in the RB TnSeq experiment (**Fig. 1**), we built a *B. ovis* strain harboring an in-frame deletion of *cysE* (Δ*cysE*). We grew Δ*cysE* in parallel with wild-type *B. ovis* ATCC 25840 (WT, **Fig. 2B**) and observed that Δ*cysE* enters stationary phase earlier and terminates growth at a lower density than WT, thus corroborating the TnSeq result. This phenotype was complemented by the addition of 4 mM cysteine to the medium (**Fig. 2B**) or by ectopic expression of the *cysE* gene from a *lac* promoter in the presence of IPTG (**Fig. 2C**). Ectopic overexpression of *cysE* in a WT *B. ovis* background did not modify growth kinetics or the growth curve shape compared to an empty vector control (WT/pSRK-EV) (**Fig. 2C**). We conclude that *cysE* and cysteine biosynthesis are necessary for normal *B. ovis* growth yield in Brucella Broth.

**Figure 2.**
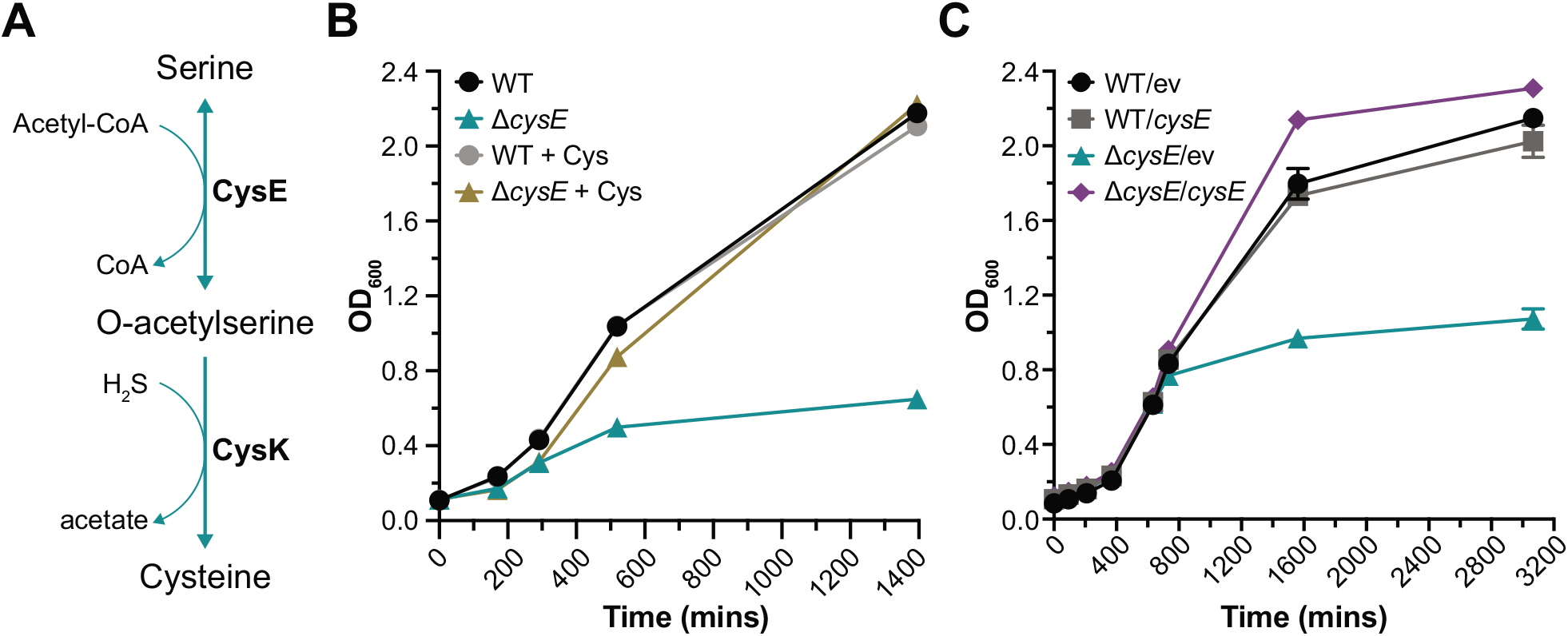
Δ*cysE* enters stationary phase prematurely; this growth defect is rescued by addition of cysteine to the growth medium. **A)** Schematic of cysteine biosynthesis from serine. Teal arrows indicate cysteine biosynthesis enzymes annotated in the *Brucella ovis* genome (RefSeq accessions NC_009505 and NC_009504). Enzymes: CysE (*BOV_RS06060, cysE*, O-acetylserine transferase); CysK1 (*BOV_RS09280, cysK*, cysteine synthase A). **B)** Representative growth curves of wild type (WT, circles), and Δ*cysE* (triangles) with (gray and ochre, respectively) or without (black and teal, respectively) addition of 4 mM cysteine (Cys) to the growth medium. **C)** Representative growth curves of WT carrying the pSRK empty vector (EV, black circle) or pSRK-*cysE* (gray square), and Δ*cysE* carrying pSRK (teal triangle) or pSRK-*cysE* (purple diamond) in BB with 1 mM IPTG, 50 μg/ml Kan. Error bars represent standard deviation of technical replicates in representative experiments.

### *cysK1* and *cysK2* function redundantly in cysteine biosynthesis

CysK catalyzes the step in cysteine biosynthesis subsequent to CysE, namely the elimination reaction in which the acetyl group on O-acetylserine is displaced by sulfide to form cysteine (20) (**Fig. 2A**). Given the stationary phase phenotype of Δ*cysE*, and the fact that this defect was chemically complemented by cysteine, we expected that mutations in cysteine synthase (*cysK*) should phenocopy Δ*cysE*. However, strains with transposon insertions in locus *BOV_RS09280* (old locus tag *BOV_1893*), annotated *cysK* in the NCBI RefSeq database, grew like wild type (**Data Set 1**). Growth of a strain harboring an in-frame deletion of *BOV_RS09280* also grew the same as WT *B. ovis* in Brucella Broth (**Fig. 3A**), confirming the TnSeq result.

**Figure 3.**
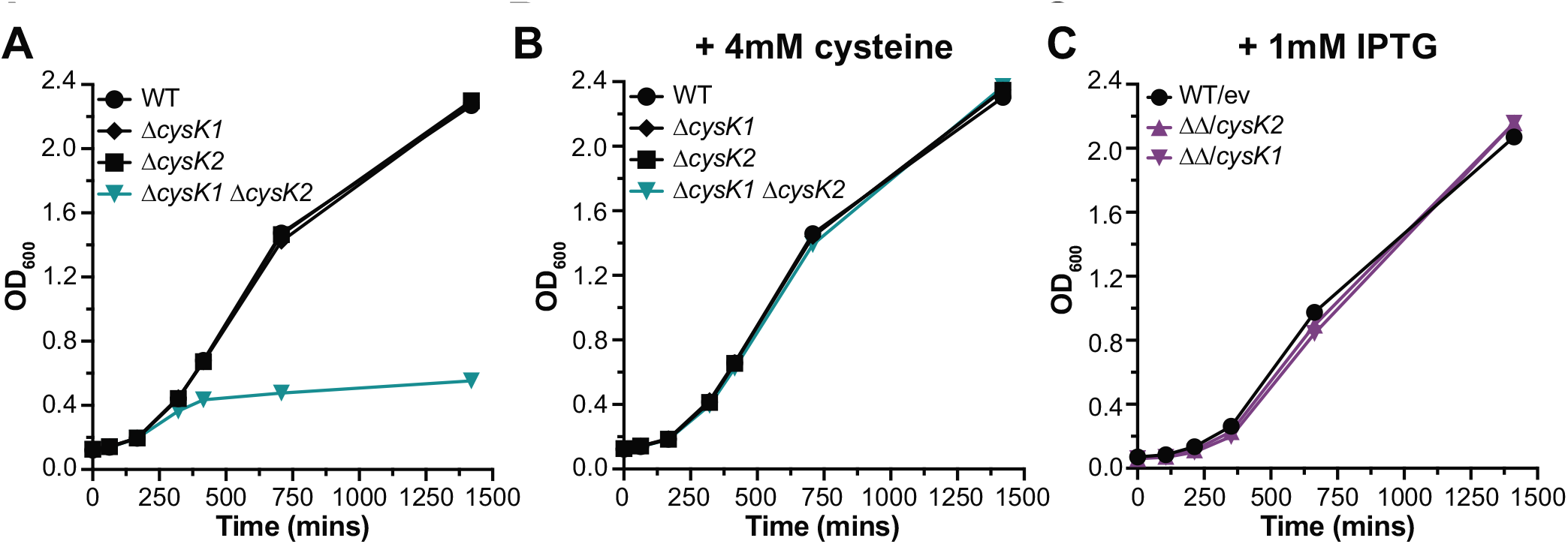
The stationary phase phenotype of a Δ*cysK1* Δ*cysK2*(*BOV_RS05050*) double deletion phenocopies Δ*cysE* and is rescued by cysteine. A)Growth of wild type (black circles), *ΔcysK1* (black diamonds), *ΔcysK2* (black squares) and *ΔcysK1 ΔcysK2* (teal triangles) in BB. B) Growth of the same four strains in A with 4 mM cysteine added to the broth. C) Growth curves of WT carrying the pSRK empty vector (black circles), and *ΔcysK1 ΔcysK2* carrying either pSRK-cysK1 or pSRK-cysK2 (purple triangles) grown with 50 μg/ml Kan and 1 mM IPTG. Error bars represent standard deviation of technical replicates and may be smaller than symbol size. Growth curves were conducted at least three independent times. A representative curve is shown for each.

We considered that the lack of an apparent growth defect in the Δ*cysK* strain may be due to the presence of other genes with CysK activity. A possible candidate for such a gene is locus *BOV_RS05050* (old locus tag *BOV_1018*), which encodes a protein with 37% sequence identity and 53% similarity to *BOV_RS09280*. Hereafter, we refer to *BOV_RS09280* as *cysK1* and *BOV_RS05050* as *cysK2*. Strains with transposon insertions in *cysK2* also yielded a wild type phenotype in our TnSeq experiment **(Data Set 1)** and a strain harboring an in-frame deletion of *cysK2* (Δ*cysK2*) likewise grew the same as WT *B. ovis*. However, a Δ*cysK1* Δ*cysK2* double deletion strain exhibited a stationary phase/growth yield phenotype similar to Δ*cysE* **(Fig. 3A)**. Like Δ*cysE*, the Δ*cysK1* Δ*cysK2* phenotype was chemically complemented by the addition of cysteine to the medium **(Fig. 3B**). The growth defect of the Δ*cysK1* Δ*cysK2* strain was genetically complemented by expressing either *cysK1* or *cysK2* from a plasmid **(Fig. 3C)**. We conclude that these two genes have redundant function in the cysteine biosynthesis pathway.

### *B. ovis* Δ*cysE* and Δ*cysK1* Δc*ysK2* strains are sensitive to exogenous H_2_O_2_ stress

Cysteine is, of course, important for protein synthesis. It is also one of the three amino acids that comprise glutathione (GSH) **(Fig. 4A)**, which plays a central role in the mitigation of a variety of stressors in bacteria (21) including oxidative stress. We hypothesized that defects in cysteine biosynthesis would have consequences on GSH synthesis and sensitize cells to oxidative stress. We thus attempted to complement the Δ*cysE* growth phenotype by adding GSH to the medium **(Fig. 4B)**. GSH supplementation partially complemented the Δ*cysE* growth yield defect. GSH limitation may directly contribute to premature entry of *B. ovis* Δ*cysE* into stationary phase, or GSH addition may restore cysteine homeostasis upon GSH catabolism. Since GSH is known to be involved in decomposition of hydrogen peroxide to water (21) **(Fig. 4A)**, we assessed whether Δ*cysE* was more sensitive to H_2_O_2_ stress. We grew WT and Δ*cysE* to stationary phase, washed the cells, treated them with H_2_O_2_ for one hour in phosphate buffered saline solution, and then enumerated CFU. The Δ*cysE* strain was ≈2000 times more sensitive to H_2_O_2_ than WT, and this sensitivity was rescued by either the addition of cysteine or glutathione to the medium during growth **(Fig. 4C)**. The hydrogen peroxide sensitivity phenotype of Δ*cysE* was genetically complemented by expression of *cysE* from the *lac* promoter of the pSRK plasmid (Δ*cysE*/pSRK-*cysE*) (**Fig. 4D**). We further tested the sensitivity of the Δ*cysK1* Δ*cysK2* double deletion mutant to hydrogen peroxide treatment. Like Δ*cysE*, the Δ*cysK1* Δ*cysK2* strain was highly sensitive to peroxide treatment. The Δ*cysK1* Δ*cysK2* peroxide survival phenotype was chemically complemented by the addition of cysteine or glutathione to the medium (**Fig. 4E**), and was genetically complemented by ectopic expression of either *cysK1* or *cysK2* (**Fig. 4F**).

**Figure 4.**
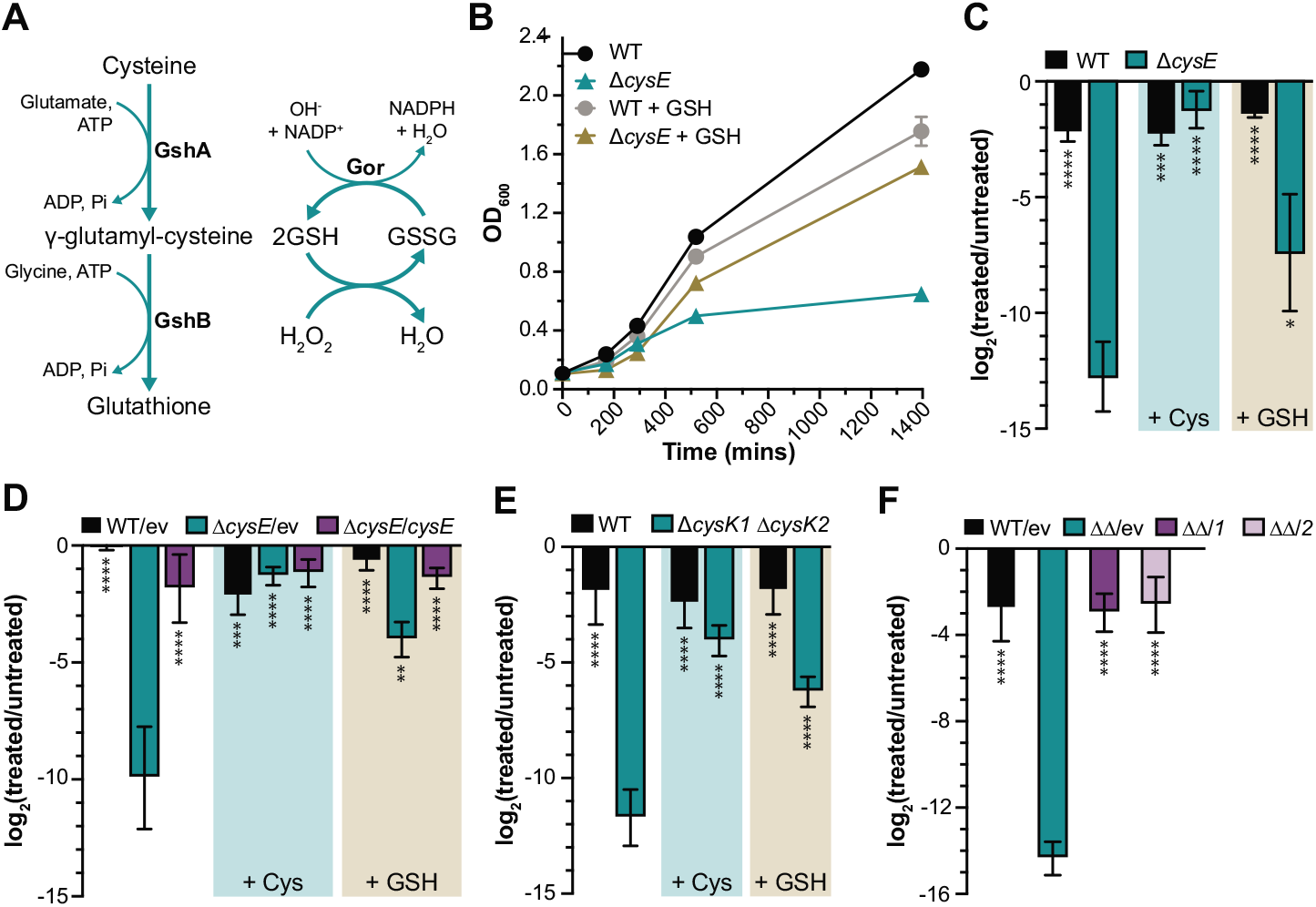
*B. ovis* Δ*cysE* is sensitive to H_2_O_2_treatment; Δ*cysE* growth defect and peroxide sensitivity is mitigated by glutathione. **A)** Schematic of glutathione metabolism. Enzymes annotated in the *Brucella ovis* genome (RefSeq accessions NC_009505 and NC_009504) are indicated in bold: GshA (*BOV_RS13935*, glutamate—cysteine ligase), GshB (*BOV_RS10075*, glutathione synthase), and Gor (*BOV_RS04850*, glutathione disulfide–reductase). GSH (glutathione, reduced state); GSSH (glutathione disulfide, oxidized state). **B)** Growth of wild type (circles) or Δ*cysE* (triangles) in BB with (gray and ochre, respectively) or without (black and teal, respectively) 4 mM GSH added to the medium. Error bars represent standard deviations and may be smaller than symbol size. **C)** Hydrogen peroxide survival assay showing the log_2_ ratio of CFU of treated (20 mM H_2_O_2_) versus untreated (mock PBS) cultures. Black bars represent wild type and teal bars represent Δ*cysE* strains. Addition of either 4 mM cysteine (+ Cys) or 4 mM GSH (+ GSH) to the medium is indicated by the shaded boxes. GSH and cysteine was washed away from the culture prior to peroxide treatment. **D)** Same assay as in **C** but with strains carrying plasmids to test genetic complementation. Strains were treated with 15 mM H_2_O_2_. Black bars represent WT carrying an empty vector (WT/ pSRK-EV), teal bars represent Δ*cysE* carrying an empty vector (Δ*cysE* / pSRK-EV), and purple bars represent complemented Δ*cysE* (Δ*cysE* / pSRK-*cysE*). p-values: * p < 0.05; ** p < 0.01; *** p < 0.001, **** p < 0.0001, calculated using one-way ANOVA (followed by Dunnett’s multiple comparison test, to Δ*cysE* in **C** or Δ*cysE*/ev in **D**). Hydrogen peroxide survival assay as in **C**, comparing wild type *B. ovis* (black bars) to the Δ*cysK1* Δ*cysK2* double deletion strain (teal bars) **F)** Same as in **D** but with strains carrying plasmids to test genetic complementation. Black bars represent *B. ovis* carrying the empty vector pSRK-EV (WT/EV), teal bars represent Δ*cysK1* Δ*cysK1* with pSRK-EV (ΔΔ/*ev*); and purple bars represent the Δ*cysK1* Δ*cysK2* strain carrying either pSRK-*cysK1* (ΔΔ/*1*, dark purple) or pSRK-*cysK2* (ΔΔ/*2*, light purple). Error bars represent standard error of the mean for 3 or 4 independent experiments.

### Δ*cysE* and *ΔcysK1 ΔcysK2* have reduced viability in the intracellular niche

*Brucella* spp. primarily reside inside mammalian host cells. There are many challenges to growth and survival in the intracellular niche including nutrient limitation and exposure to stressors such as reactive oxygen species (ROS) (6). Given the *in vitro* growth and hydrogen peroxide sensitivity phenotypes of the *cys* mutants, we tested whether fitness of the Δ*cysE* and Δ*cysK1* Δ*cysK2* strains was compromised in the intracellular niche. We infected a human monocytic cell line, THP-1, that we differentiated into macrophage-like cells. Although entry (2 hrs post-infection, p.i.) to the macrophage was unaffected by deletion of *cysE* or *cysK1* and *cysK2*, there was a significant loss in recoverable colony forming units (CFU) of the *cys* mutant strains by 24 hrs p.i. relative to WT **(Fig. 5A and 5B)**. Thus *B. ovis* Δ*cysE* and Δ*cysK1* Δ*cysK2* enter host cells like WT but their fitness is compromised after entry.

**Figure 5.**
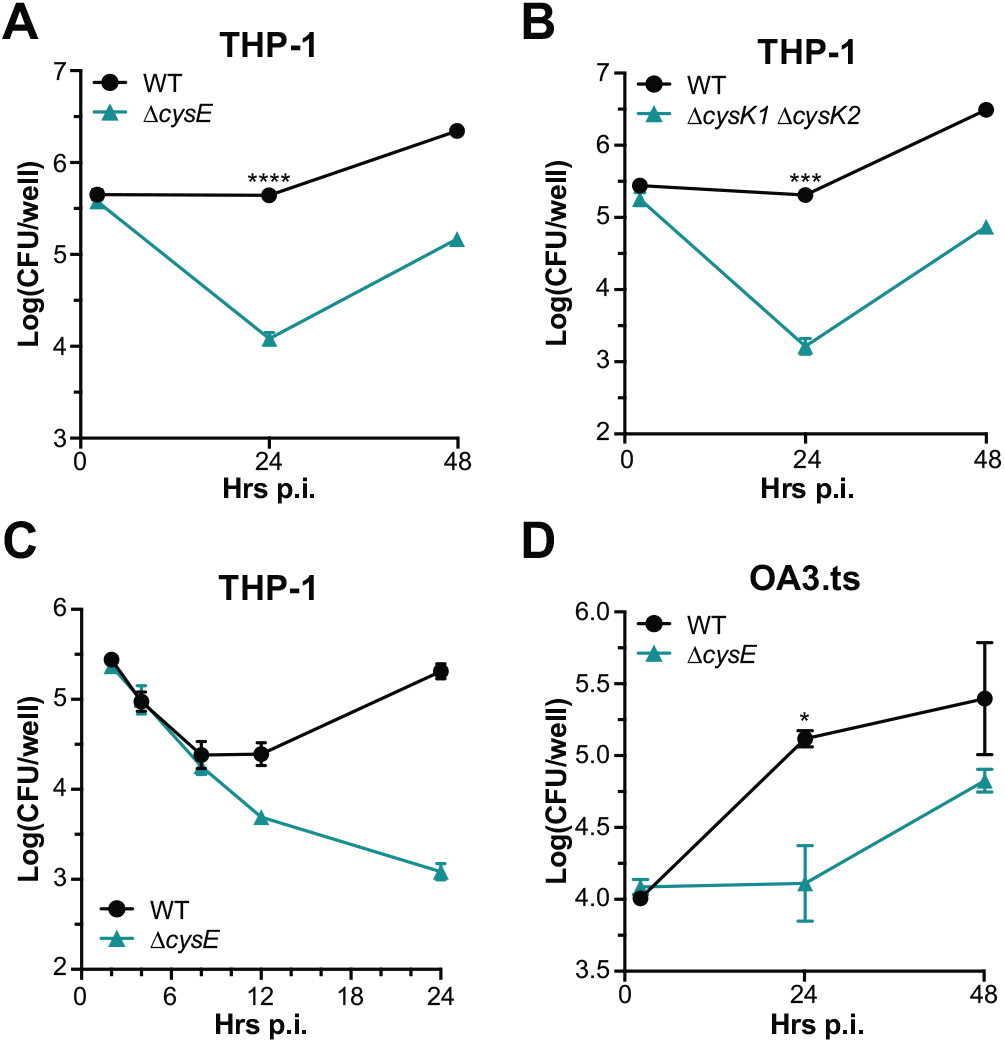
*B. ovis* Δ*cysE* has reduced fitness in the intracellular niche of human macrophage-like cells and an ovine testis epithelial cell line. Log_10_ colony forming units (CFU) per well of WT (black circles) or Δ*cysE* (teal triangles) isolated from infected THP-1 (**A** and **C**) or OA3.ts mammalian cells (**D**) enumerated at 2, 24 and 48 hours (hrs) post infection (p.i.) (**A** and **D**) or at 2, 4, 8, 12, and 24 hrs p.i. (**C**). **B)** Log_10_ CFU/well per well of WT (black circles) or Δ*cysK1* Δ*cysK2* (teal triangles) strains isolated from infected THP-1 enumerated at 2, 24 and 48 hrs p.i. p-value comparing recovered *B. ovis* 24 hrs p.i. were calculated using an unpaired t-test **A, B** and **D**. * p < 0.05; *** p < 0.001, **** p < 0.0001. Infections were repeated 3-5 independent times. Error bars represent standard error within the representative experiment.

In an effort to distinguish the relative contributions of intracellular killing and cysteine (nutritional) limitation on reduced fitness of the *cys* mutants, we enumerated CFU recovered from THP-1 cells at timepoints between 2 and 24 hrs p.i. Wild type and Δ*cysE* exhibit identical CFU loss between 2 and 8 hrs post-infection. WT begins to replicate by 12 hrs, but Δ*cysE* CFUs continue to decrease up to 24 hrs **(Fig. 5C)**. The rate of recoverable CFU increase between 24 hrs and 48 hrs p.i. is similar between WT and the *cys* mutants providing evidence that *B. ovis* has access to cysteine in this environment (Fig. 5A and 5B). An attempt to interrogate a later time point (72 hrs) was confounded by egressing bacteria and subsequent killing by gentamicin in the tissue culture medium **(Fig. S3)**. The intracellular infection defect we observe was complemented by expression of *cysE* from a plasmid **(Fig. S4A)**. We attribute partial complementation to the fact that *cysE* was expressed from a *lac* promoter on a replicating plasmid; there are challenges with full induction of transgenes from heterologous promoters in an intracellular infection context.

Given the ability of *Brucella* to infect multiple mammalian cell types, we next tested whether the *in vitro* infection phenotype of the *cys* mutants was particular to macrophages. We infected a sheep testis epithelial cell line (OA3.ts) (22), which is derived from a host tissue type that is relevant to *B. ovis* infection. OA3.ts entry was unaffected by the lack of *cysE*, but recovered CFU were significantly lower for Δ*cysE* by 24 hr p.i. **(Fig. 5D)**. This phenotype was partially complemented by ectopic expression of *cysE* from a plasmid **(Fig. S4B)**. The magnitude of the Δ*cysE* defect 24 hrs p.i. was greater in THP-1 macrophages than in the OA3.ts epithelial line (about 64-fold vs 16-fold, respectively; Fig. S5). These *in vitro* infection data provide evidence that an intact cysteine metabolism system promotes *B. ovis* fitness in intracellular niche of multiple mammalian cell types.

## Discussion

*A genome-scale search for B. ovis stationary phase mutants leads to cysteine metabolism*

*B. ovis* is a widespread ovine pathogen that remains an understudied member of the *Brucella* genus. Using a RB TnSeq approach, we sought to identify genes that are important for *B. ovis* growth and/or survival in the late phase of axenic broth culture (i.e. stationary phase), with a larger goal of uncovering genes that are important for fitness in the intracellular environment. We identified multiple genes for which Tn-*himar* disruption resulted in reduced fitness in stationary phase. Among the expected mutants in this dataset is *rsh* (*BOV_RS03230*), which controls the stringent response (23). Additionally, genes involved in purine metabolism, including *purF* have reduced fitness in dense culture. In *Mycobacterium smegmatis*, PurF influences survival during stationary phase (24), and purine metabolism is known to be important for growth of multiple microbes in the intracellular and extracellular environments (25, 26). Multiple genes with a predicted role in tRNA modification also had diminished fitness in stationary phase. Transfer RNA modification enzymes have roles in translation quality control and can function to direct translation of specific transcripts under particular growth conditions (27). Given the phenotypes of tRNA modification mutants in stationary phase, it may be the case that these genes play a role of regulation of *Brucella ovis* physiology in the intracellular niche.

Tn-*himar* strains with insertions in *cysE* had the most diminished fitness in stationary phase, and *cysE* was therefore selected for follow-up studies. Sulfur and cysteine metabolism are central to microbial growth, and have been well studied in numerous pathogens (28). In *Brucella* spp., our understanding of sulfur metabolism is relatively limited though biosynthesis of sulfur-containing amino acids - cysteine and methionine - has been implicated in *Brucella melitensis* 16M infection of mice (29). Based on the high level of sequence identity/similarity to well-studied CysE enzymes and established structural data on *B. abortus* CysE (30), *B. ovis* CysE is presumed to catalyze biosynthesis of O-acetylserine from acetyl-CoA and serine. The subsequent step in biosynthesis of cysteine from O-acetylserine requires displacement of the acetyl group by sulfide, a reaction that is catalyzed by CysK in many bacteria. Our growth data clearly implicate *cysE* in the cysteine biosynthesis pathway, as the *in vitro* growth defect of Δ*cysE* is rescued by the addition of cysteine. These results support published data that *Brucella* spp. can assimilate cysteine as an exogenous organic sulfur source (31, 32).

Surprisingly, the growth phenotypes of strains with Tn-*himar* insertions in the gene annotated as *cysK* in the RefSeq database did not differ from wild type, which suggested redundancy at this biosynthetic step. Consistent with this hypothesis, we have presented genetic evidence that two related enzymes, CysK1 and CysK2, function redundantly to produce cysteine. It is possible that CysK1 and CysK2 do not catalyze the same reaction, but rather determine cysteine biosynthesis through two distinct routes. A recent report of such a case is the cystathionine β-synthase of *Helicobacter pylori*, which retains some O-acetylserine sulfhydrylase activity (33); this enzyme shares primary structure features with *B. ovis* CysK2. CysK-family enzymes can also have functions beyond direct involvement in cysteine metabolism (34), which may influence interpretation of our results. Notably, *B. abortus* CysE (serine O-acetyltransferase) and CysK2 do not form a cysteine synthase complex (CSC) *in vitro* (35). This supports a model in which CysK2 participates in cysteine synthesis via a mechanism that differs from that catalyzed by the typical CysE-CysK CSC. We postulate that CysK1, rather than CysK2, binds to CysE to form the CSC in *Brucella*. The development of a defined medium that supports the growth of *B. ovis* would greatly facilitate future study of CysK1 and CysK2 functions in cells. Exploration of the possible intracellular fitness advantage gained by redundancy at the CysK step of cysteine biosynthesis is an interesting area of future investigation.

### Cysteine, glutathione and hydrogen peroxide stress

The growth and peroxide survival defects of Δ*cysE* were partially rescued by addition of cysteine or glutathione to the medium. Though elevated intracellular cysteine enhances susceptibility to hydrogen peroxide stress in *Escherichia coli* (36, 37), we do not observe peroxide sensitization of WT or Δ*cysE B. ovis* upon addition of 4 mM cysteine. 4 mM cysteine was consistently more protective than 4 mM GSH in our assay. GSH is an important redox control molecule, but the protective effect of GSH supplementation against H_2_O_2_ may be indirect. Specifically, it’s possible that *B. ovis* transports and metabolizes some of the GSH to release cysteine, which is one of the three component amino acids of GSH. *B. ovis* is predicted to encode a γ-glutamylcyclotransferase (*BOV_RS09395*), which catalyzes the cleavage of GSH to form pyroglutamic acid and L-cysteinylglycine (38). The L-cysteinylglycine dipeptide could then be separated by peptidases to release cysteine. We nonetheless favor a model in which diminished GSH production (as a result of abolished cysteine production) in Δ*cysE* directly affects H_2_O_2_ detoxification and growth yield of Δ*cysE*. Glutathione metabolism is important in *B. ovis*: the *gshA* biosynthesis gene is essential based on our previously published TnSeq dataset (16). Moreover, Tn-*himar* insertions in *BOV_RS04850* (old locus tag, *BOV_0978*), which is predicted to encode a glutathione-disulfide reductase (*gor*) – that reduces GSSG to GSH – resulted in a significant fitness disadvantage throughout the growth curve.

### Cysteine and growth and the intracellular niche

Our study provides evidence that cysteine biosynthesis contributes to *B. ovis* fitness inside mammalian host cells. Strains harboring deletions of *cysE* or both *cysK1* and *cysK2* were not defective in host cell entry but had significantly reduced recoverable CFUs at 24 and 48 hrs post infection in human macrophage-like and ovine epithelial cell lines. Reduced recoverable CFU of the *cys* mutants at 24 and 48 hours can be attributed to defects that are manifested between 2 and 24 hours post-infection. It is difficult to fully discern the relative contributions of nutritional restriction and enhanced oxidative stress sensitivity to attenuation of Δ*cysE* and Δ*cysK1* Δ*cysK2 in vitro*. The similar observed rate at which recoverable CFUs increase between 24 and 48 hours suggests that cysteine levels are not limiting – at least after 24 hours. The ER-derived replicative *Brucella* containing vacuole (rBCV) supports bacterial replication (39) and can be established as early as 12 hrs p.i. (40); based on our data, we conclude that this compartment contains enough cysteine or cysteine-containing peptides to support growth. Of note, the defect of the Δ*cysE* strain was more pronounced in macrophage-like cells than in an ovine testis epithelial line. Sensitivity of Δ*cysE* and Δ*cysK1* Δ*cysK2* to ROS may underlie this difference in fitness between cell lines as macrophages typically have a more robust respiratory burst than epithelial cells (41). A significant decrease in recoverable CFUs in Δ*cysE* is evident by 4 hr post-infection. This is a timepoint before intracellular *Brucella* replication occurs (40), supporting a model in which increased sensitivity to host killing underlies the reduced fitness of Δ*cysE* and Δ*cysK1* Δ*cysK2* in host cells.

Previous *B. abortus* TnSeq studies by Sternon and colleagues (42) did not identify *cysE* as a gene that was important for infection of Raw 264.7 macrophages. We observed significantly reduced *B. ovis* Δ*cysE* CFU relative to WT in Raw 264.7 cells by 24 hrs **(Fig. S6)**, but the Sternon *et al*. experiment and our experiment differ in several ways. Indeed, the importance of cysteine metabolism in intracellular growth and/or survival may vary between *B. ovis* and *B. abortus* and between mammalian cell lines. Nonetheless, our data clearly provide evidence that a cysteine anabolism pathway in *B. ovis* is important for growth, stress survival and fitness in the intracellular niche.

Cysteine and methionine metabolic pathways are attractive targets to combat various pathogens (28) because, in mammals, these amino acids must be acquired from diet. Thus, compounds that disrupt cysteine metabolism are not predicted to have direct negative effects on mammalian metabolism. In fact, O-acetylserine sulfhydrylase (OASS; i.e. *cysK*) inhibitors are under investigation as therapeutics for *Mycobacterium tuberculosis* infections (43). Our work shows that genetic disruption of cysteine biosynthesis leads to a significant defect in *B. ovis* fitness within host cells. This pathway is therefore a possible target for combating brucellosis.

## Materials and Methods

### Bacterial strains and growth conditions

*Brucella ovis* was grown on Tryptic Soy Agar (TSA, Difco Laboratories) plates, supplemented with 5% sheep blood (Quad Five) or in Brucella Broth (BB, Difco Laboratories, dissolved in tap water) for liquid cultures. Cells were incubated at 37 °C with 5% CO_2_ supplementation. Kanamycin (Kan) 50 μg/ml, sucrose (5% w/v) or isopropyl β-D-1-thiogalactopyranoside (IPTG, GoldBio) at 1 mM or 2 mM, were added when required.

*Escherichia coli* strains were grown in lysogeny broth (LB, Fisher Bioreagents) or on LB + 1.5% agar (Fisher Bioreagents) plates at 37 °C with Kan supplemented at a concentration of 50 μg/ml when required. *E. coli* WM3064 strain, used for conjugation, was grown in the presence of 300 μM diaminopimelic acid (DAP, Sigma-Aldrich), as it is a DAP auxotroph.

### Plasmid and strain construction

#### Deletion plasmid construction

To build the deletion strains, fragments of approximately 500 bp upstream and downstream of target genes were amplified with KOD Xtreme Hot Start polymerase (Novagen). These fragments were built so that 9 bases at both the 5’ and 3’ ends of the gene were maintained, keeping the gene product in frame to minimize polar effects. Purified DNA from *Brucella ovis* ATCC 25840 was used as a template. Amplified fragments were gel purified (ThermoFisher Scientific) and assembled into the pNPTS138 suicide deletion vector (digested with HindIII and BamHI restriction enzymes, New England Biolabs) using Gibson assembly (New England Biolabs).

#### Complementation plasmid construction

To build plasmids for genetic complementation, *cysE, cysK1* or *cysK2* were PCR amplified from *B. ovis* ATCC 25840 with KOD Xtreme Hot Start polymerase, gel purified (ThermoFisher Scientific) and Gibson assembled into pSRK (44) that had been digested with NdeI and KpnI restriction enzymes (New England Biolabs). *cysE, cysK1* or *cysK2* were cloned downstream of P_lac_ (lactose, IPTG inducible promoter).

#### Delivery of plasmids to B. ovis

Constructed plasmids were transformed into chemically competent *E. coli* Top10 strains for plasmid maintenance. All plasmid inserts were confirmed by PCR and Sanger sequencing, and plasmids were delivered to *B. ovis* by conjugation using *E. coli* WM3064 as a donor strain. For conjugation, WM3064 donor strains were mated with *B. ovis* strains and spotted on TSA blood plates plus DAP and incubated overnight at 37° C in a 5% CO_2_ atmosphere. Mating spots were spread on TSA blood plates plus Kan (without DAP) to select for *B. ovis* plasmid acquisition. When deleting genes using the pNPTS138 plasmid, merodiploid clones were inoculated in Brucella Broth overnight to allow for a second crossover event, then spread on TSA blood plates plus sucrose (5% w/v) for counterselection. Single colonies harboring gene deletions were identified by patching clones on TSA blood plates with or without Kan. The putative deleted locus was PCR amplified using gene-flanking primers in Kan-sensitive clones, and the PCR fragment was resolved by gel electrophoresis to test whether the gene had been deleted. For a complete list of strains, plasmids, and primers, please see **Data Set 3**.

### Growth Curves

Cells were inoculated from ∼48hr-old TSA blood plates into BB at densities ranging from OD_600_0.08 to OD_600_ of 0.2. Growth was assessed spectrophotometrically measuring optical density at 600 nm (OD_600_). Growth curves were conducted at least three independent times with two or three technical replicated in each experiment. Representative curves are shown for each set of strains. Where indicated, cysteine (4 mM), GSH (4 mM), Kan (50 μg/μl), or IPTG (1 mM) were supplemented upon start of growth experiment to the liquid media.

### H_2_O_2_ survival assays

Cells were grown overnight in BB to stationary phase (OD_600_ of ≈ 2). Cells were pelleted and resuspended in Phosphate-Buffered Saline (PBS, Sigma) to achieve an OD_600_ of 0.15. 200 μl of cells were added to 1.8 ml of PBS or PBS supplemented with fresh H_2_O_2_ (15 or 20 mM final concentration), bringing the final OD_600_ to 0.015. Cells were then incubated at 37 °C in 5% CO_2_ for 1 hr before spotting aliquots of a 10-fold serial dilution series on TSA blood plates. CFUs were enumerated after 48 hrs incubation at 37 °C in 5% CO_2_. Experiments were repeated at least three times with each sample in duplicate or triplicate in each experiment.

### DNA extractions

Cells from 1 ml of stationary phase culture were pelleted by centrifugation, washed once in PBS, and resuspended in 100 μl TE buffer (10 mM Tris-HCl, 1 mM EDTA, pH 8.0) supplemented with 1 μg/ml RNAseA. Cells were lysed by addition of 0.5 ml GES lysis solution (5 M guanidinium thiocyanate, 0.5 M EDTA pH 8.0, 0.5% v/v Sarkosyl) and 15 min incubation at 60 °C. 0.25 ml cold 7.5 M ammonium acetate (Fisher Bioreagents) was added, and mixture was incubated on ice for 10 min. 0.5 ml of chloroform (Fisher Bioreagents) was added to separate the DNA, samples were vortexed and centrifuged. Aqueous top phase was moved to a fresh 1.5 ml centrifuge tube and 0.54 volumes of cold isopropanol was added to precipitate the DNA. After centrifugation, isopropanol was discarded and pellets were washed three times in 70% ethanol before resuspending pellets in TE buffer. Concentration and purity of the extracted DNA were determined spectrophotometrically (NanoDrop One, Thermo Scientific).

### Barcoded TnSeq

A *B. ovis* RB Tn-*himar* library was built and mapped as described (31). Briefly, *E. coli* APA752 (a WM3064 donor strain carrying a pKMW3 *mariner* transposon library) was conjugated into *B. ovis bcaA1* (16) under atmospheric CO_2_ conditions. Kan resistant colonies were collected, grown to OD_600_ = 0.6 and frozen in 1 ml aliquots. Genomic DNA was extracted and the Tn insertion sites mapped as previously described (19).

To identify genes that confer a fitness advantage in stationary phase, the *B. ovis* Tn-*himar* library was inoculated in BB in a 5% CO_2_ environment in triplicate at an OD_600_ of 0.0025. An aliquot of each initial culture was collected as the reference time point. Cultures were then grown to stationary phase, with samples harvested throughout the growth curve at OD_600_ of 0.05, 0.12, 0.9 and 2.4. Cells from each sample were pelleted by centrifugation and resuspended in water. Barcodes were amplified from approximately 1.5 × 10^8^ cells per PCR reaction (see **Table S1**) with primers that both amplified the barcodes and added indexed adaptors (19). Amplified barcodes were then pooled, purified and sequenced on an Illumina HiSeq 4000.

Fitness scores for each gene were calculated following the protocol of Wetmore and colleagues (19) using scripts available at https://bitbucket.org/berkeleylab/feba. Genes for which 2 out of 3 samples had at least one timepoint with a t-like statistical significance score (19) ≥ |4| were included in subsequent analyses. A heat map of fitness scores of genes passing this filter is shown in **Fig. S1** and the raw fitness data are in **Data Set 1**.

Finally, we averaged the fitness values of the three replicates and kept mutants that an average fitness score ≥ |1| in at least one time point. Mutants in this group with a standard deviation ≥ 0.75, were manually inspected and extreme outlier points were removed from a total of six genes. The genes with adjusted average and standard deviation values are shown in red in **Data Set 1**. Heat map of averaged fitness values is shown in **Fig. 1**.

### Tissue culture

All tissue culture cells were grown at 37°C with 5% CO_2_ supplementation. THP-1 cells (ATCC TIB-202) were cultured in Roswell Park Memoriam Institute medium (RPMI 1640, Gibco) + 10% Fetal Bovine Serum (FBS, Fisher Scientific). The RAW 264.6 (ATCC TIB-71) and the OA3.ts (ATCC CRL-6546) cells were grown in Dulbecco’s Modified Eagle Medium (DMEM, Gibco) supplemented with 10% FBS.

### Infection assays

THP-1 cells were seeded at a concentration of 10^5^ cells/well in 96-well plates and phorbol myristate acetate (PMA) at final concentration of 50 ng/μl was added to induce differentiation into macrophage-like cells for 48-96 hrs prior to infection. OA3.ts cells were seeded at a density of 5 × 10^4^ cells/well in 96-well plates for 24 hrs prior to infection. *B. ovis* cells were resuspended from a 48 hr old plate in RPMI + 10% FBS or DMEM + 10% FBS and added to tissue culture plates on the day of infection at multiplicity of infection (MOI) of 100 for THP-1, and at an MOI of 1000 for OA3.ts cells. When infecting with the complementation strains carrying the pSRK plasmid, the *Brucella* strains were struck on TSA blood plates with Kan and IPTG 48 hrs prior to infection, and 2 mM IPTG was added to the tissue culture media throughout the duration of the experiment. Plates were spun for 5 min at 150 × g and incubated for 1 hr at 37 °C in 5% CO_2_. Fresh media was supplied containing 50 μg/ml of gentamicin and incubated for another hour. Cells were then washed once with PBS and once in H_2_O and then lysed with H_2_O for 10 min RT at 2 hrs, 24 hrs, and 48 hrs post infection. Lysates were serially diluted, spotted on TSA blood plates and incubated at 37 °C in 5% CO_2_ for 48 hrs to enumerate CFUs. Experiments were repeated at least 3 times with three technical replicates.

## Acknowledgements

We thank David Hershey for providing critical constructive feedback and help with the RB Tn-*himar* library experiment, as well as Crosson lab members, Brendan MacNabb, and Josh Lensmire for helpful discussions. This work was funded through the NIH Training Grant T32 GM007197 (L.M.V.), NIH grant R35 GM131762 (S.C.), and the Gallo Global Health Fellowship Program (L.M.V.).

## References

1. Roop II RM, Gee JM, Robertson GT, Richardson JM, Ng W-L, Winkler ME. 2003. Brucella stationary-phase gene expression and virulence. Annu Rev Microbiol 57:57–76.

2. Porte F, Liautard JP, Köhler S. 1999. Early acidification of phagosomes containing Brucella suis is essential for intracellular survival in murine macrophages. Infect Immun 67:4041– 4047.

3. Jiang X, Leonard B, Benson R, Baldwin CL. 1993. Macrophage control of Brucella abortus: Role of reactive oxygen intermediates and nitric oxide. Cell Immunol 151:309–319.

4. Brundu S FA. 2015. Polarization and repolarization of macrophages. J Clin Cell Immunol 06:1–10.

5. Muraille E, Leo O, Moser M. 2014. Th1/Th2 paradigm extended: Macrophage polarization as an unappreciated pathogen-driven escape mechanism? Front Immunol 5:1–12.

6. Roop RM, Gaines JM, Anderson ES, Caswell CC, Martin DW. 2009. Survival of the fittest: How Brucella strains adapt to their intracellular niche in the host. Med Microbiol Immunol 198:221– 238.

7. Kim HS, Caswell CC, Foreman R, Roop RM, Crosson S. 2013. The Brucella abortus general stress response system regulates chronic mammalian infection and is controlled by phosphorylation and proteolysis. J Biol Chem 288:13906–13916.

8. Kim HS, Willett JW, Jain-Gupta N, Fiebig A, Crosson S. 2014. The Brucella abortus virulence regulator, LovhK, is a sensor kinase in the general stress response signalling pathway. Mol Microbiol 94:913–925.

9. Rossetti CA, Galindo CL, Lawhon SD, Garner HR, Adams LG. 2009. Brucella melitensis global gene expression study provides novel information on growth phase-specific gene regulation with potential insights for understanding Brucella:host initial interactions. BMC Microbiol 9:1–14.

10. Buddle M, Boyes B. 1953. A Brucella mutant causing genital disease of sheep in New Zealand. Aust Vet J 29:145–153.

11. Tsolis RM, Seshadri R, Santos RL, Sangari FJ, JM García Lobo, de Jong MF, Ren Q, Myers G, Brinkac LM, Nelson WC, DeBoy RT, Angiuoli S, Khouri H, Dimitrov G, Robinson JR, Mulligan S, Walker RL, Elzer PE, Hassan KA, Paulsen IT. 2009. Genome degradation in Brucella ovis corresponds with narrowing of its host range and tissue tropism. PLoS One https://doi:10.1371/journal.pone.0005519.

12. Wattam AR, Foster JT, Mane SP, Beckstrom-Sternberg SM, Beckstrom-Sternberg JM, Dickerman AW, Keim P, Pearson T, Shukla M, Ward D V., Williams KP, Sobral BW, Tsolis RM, Whatmore AM, O’Callaghan D. 2014. Comparative phylogenomics and evolution of the brucellae reveal a path to virulence. J Bacteriol 196:920–930.

13. Antunes JMA de P, Allendorf SD, Appolinário CM, Cagnini DQ, Figueiredo PR, Júnior JB, Baños JV, Kocan KM, de la Fuente J, Megid J. 2013. Rough virulent strain of Brucella ovis induces pro-and anti-inflammatory cytokines in reproductive tissues in experimentally infected rams. Vet Microbiol 161:339–343.

14. Gouletsou PG, Fthenakis GC. 2015. Microbial diseases of the genital system of rams or bucks. Vet Microbiol 181:130–135.

15. Picard-Hagen N, Berthelot X, Champion JL, Eon L, Lyazrhi F, Marois M, Peglion M, Schuster A, Trouche C, Garin-Bastuji B. 2015. Contagious epididymitis due to Brucella ovis: Relationship between sexual function, serology and bacterial shedding in semen. BMC Vet Res 11:1–7.

16. Varesio LM, Willett JW, Fiebig A, Crosson S. 2019. A carbonic anhydrase pseudogene sensitizes select Brucella lineages to low CO2 tension. J Bacteriol https://doi.org/10.1128/JB.00509-19.

17. MacHelart A, Willemart K, Zúñiga-Ripa A, Godard T, Plovier H, Wittmann C, Moriyón I, De Bolle X, Van Schaftingen E, Letesson JJ, Barbier T. 2020. Convergent evolution of zoonotic Brucella species toward the selective use of the pentose phosphate pathway. Proc Natl Acad Sci U S A 117:26374–26381.

18. Ronneau S, Moussa S, Barbier T, Conde-Álvarez R, Zuniga-Ripa A, Moriyon I, Letesson JJ. 2016. Brucella, nitrogen and virulence. Crit Rev Microbiol 42:507–525.

19. Wetmore KM, Price MN, Waters RJ, Lamson JS, He J, Hoover CA, Blow MJ, Bristow J, Butland G, Arkin AP, Deutschbauer A. 2015. Rapid quantification of mutant fitness in diverse bacteria by sequencing randomly bar-coded transposons. MBio 6:1–15.

20. Kredich NM, Tomkins GM. 1966. The enzymic synthesis of L-cysteine in Escherichia coli and Salmonella typhimurium. J Biol Chem 241:4955– 4965.

21. Masip Ll, Veeravalli K, Georgiou G. 2006. The many faces of glutathione in bacteria. Antioxidants Redox Signal 8:753–762.

22. Sidhu-Muñoz RS, Sancho P, Vizcaíno N. 2018. Evaluation of human trophoblasts and ovine testis cell lines for the study of the intracellular pathogen Brucella ovis. FEMS Microbiol Lett 365:1–9.

23. Dozot M, Boigegrain RA, Delrue RM, Hallez R, Ouahrani-Bettache S, Danese I, Letesson JJ, De Bolle X, Köhler S. 2006. The stringent response mediator Rsh is required for Brucella melitensis and Brucella suis virulence, and for expression of the type IV secretion system virB. Cell Microbiol 8:1791–1802.

24. Keer J, Smeulders MJ, Williams HD. 2001. A purF mutant of Mycobacterium smegmatis has impaired survival during oxygen-starved stationary phase. Microbiology 147:473–481.

25. Samant S, Lee H, Ghassemi M, Chen J, Cook JL, Mankin AS, Neyfakh AA. 2008. Nucleotide biosynthesis is critical for growth of bacteria in human blood. PLoS Pathog doi:10.1371/journal.ppat.0040037.

26. Shaffer C, Guckes KR, Breland EJ, Floyd KA, Casella DP, Algood HMS, Clayton DB. 2017. Purine biosynthesis metabolically constrains intracellular survival of uropathogenic Escherichia coli. Infect Immun 85:1–14.

27. de Crécy-Lagard V, Jaroch M. 2020. Functions of bacterial tRNA modifications: from ubiquity to diversity. Trends Microbiol https://doi.org/10.1016/j.tim.2020.06.010.

28. Lensmire JM, Hammer ND. 2019. Nutrient sulfur acquisition strategies employed by bacterial pathogens. Curr Opin Microbiol 47:52–58.

29. Lestrate P, Delrue RM, Danese I, Didembourg C, Taminiau B, Mertens P, De Bolle X, Tibor A, Tang CM, Letesson JJ. 2000. Identification and characterization of in vivo attenuated mutants of Brucella melitensis. Mol Microbiol 38:543–551.

30. Kumar S, Kumar N, Alam N, Gourinath S. 2014. Crystal structure of serine acetyl transferase from Brucella abortus and its complex with coenzyme A. Biochim Biophys Acta - Proteins Proteomics 1844:1741–1748.

31. Herrou J, Willett JW, Fiebig A, Varesio LM, Czyż DM, Cheng JX, Ultee E, Briegel A, Bigelow L, Babnigg G, Kim Y, Crosson S. 2019. Periplasmic protein EipA determines envelope stress resistance and virulence in Brucella abortus. Mol Microbiol 111:637–661.

32. Rode LJ, Lankford CE, Schuhardt VT. 1951. Studies of sulfur metabolism of Brucella suis. J Bacteriol 62:571–582.

33. Devi S, Tarique KF, Ali MF, Abdul Rehman SA, Gourinath S. 2019. Identification and characterization of Helicobacter pylori O-acetylserine-dependent cystathionine β-synthase, a distinct member of the PLP-II family. Mol Microbiol 112:718–739.

34. Campanini B, Benoni R, Bettati S, Beck CM, Hayes CS, Mozzarelli A. 2015. Moonlighting O-acetylserine sulfhydrylase: New functions for an old protein. Biochim Biophys Acta - Proteins Proteomics 1854:1184–1193.

35. Dharavath S, Raj I, Gourinath S. 2017. Structure-based mutational studies of O-acetylserine sulfhydrylase reveal the reason for the loss of cysteine synthase complex formation in Brucella abortus. Biochem J 474:1221–1239.

36. Korshunov S, Imlay KRC, Imlay JA. 2020. Cystine import is a valuable but risky process whose hazards Escherichia coli minimizes by inducing a cysteine exporter. Mol Microbiol 113:22–39.

37. Park S, Imlay JA. 2003. High levels of intracellular cysteine promote oxidative DNA damage by driving the Fenton reaction. J Bacteriol 185:1942–1950.

38. Meister A, Anderson ME. 1983. Glutathione. Annu Rev Biochem 52:711–760.

39. Celli J. 2015. The changing nature of the Brucella-containing vacuole. Cell Microbiol 17:951–958.

40. Celli J. 2019. The intracellular life cycle of Brucella spp. Microbiol Spectr doi:10.1128/microbiolspec.BAI-0006-2019.

41. Lambeth JD. 2004. NOX enzymes and the biology of reactive oxygen. Nat Rev Immunol 4:181–189.

42. Sternon JF, Godessart P, de Freitas RG, Van der Henst M, Poncin K, Francis N, Willemart K, Christen M, Christen B, Letesson JJ, De Bolle X. 2018. Transposon sequencing of Brucella abortus uncovers essential genes for growth in vitro and inside macrophages. Infect Immun 86:1–20.

43. Schnell R, Sriram D, Schneider G. 2015. Pyridoxal-phosphate dependent mycobacterial cysteine synthases: Structure, mechanism and potential as drug targets. Biochim Biophys Acta - Proteins Proteomics 1854:1175–1183.

44. Khan SR, Gaines J, Roop 2nd RM, Farrand SK, Roop RM, Farrand SK. 2008. Broad-host-range expression vectors with tightly regulated promoters and their use to examine the influence of TraR and TraM expression on Ti plasmid quorum sensing. Appl Env Microbiol 74:5053–5062.

