## Supplemental Figures for "*Brucella ovis* cysteine biosynthesis contributes to peroxide stress survival and fitness in the intracellular niche"

### Supplemental Material

##### Table S1

| Timepoint | $OD_{600}$ | Volume | Number of cells collected | Cells in each PCR rxn |
|---|---|---|---|---|
| T0 | 0.6 | 100 ml | $4 \times 10^9$ | $8 \times 10^7$ |
| T1 | 0.05 | 1 ml | $6.6 \times 10^9$ | $1.3 \times 10^8$ |
| T2 | 0.12 | 1 ml | $8 \times 10^9$ | $1.6 \times 10^8$ |
| T3 | 0.9 | 0.5 ml | $6 \times 10^9$ | $1.2 \times 10^8$ |
| T4 | 2.4 | 0.5 ml | $8 \times 10^9$ | $1.6 \times 10^8$ |

rxn = reaction;

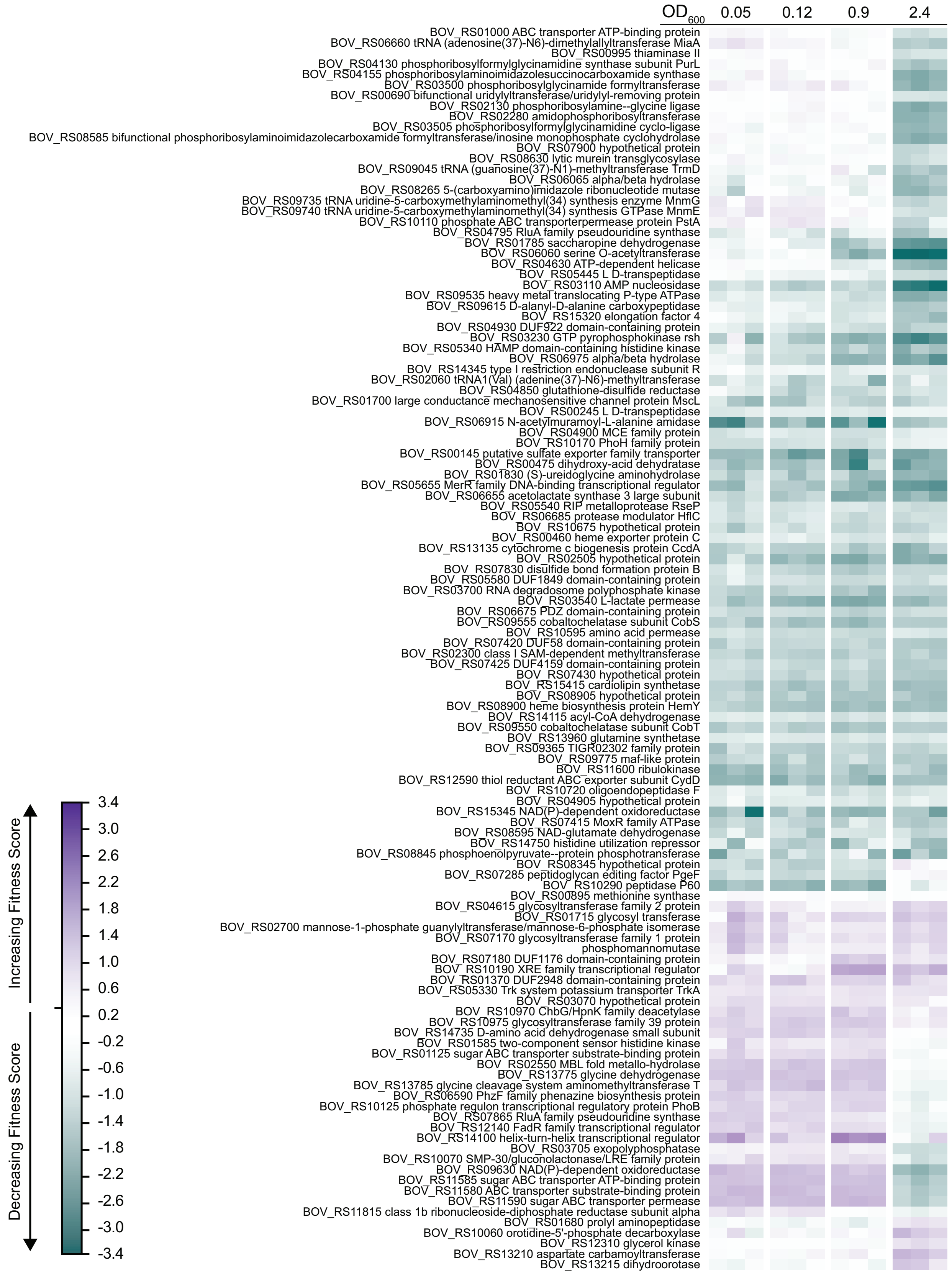
OD600 0.05 0.12 0.9 2.4
BOV_RS01000 ABC transporter ATP-binding protein
BOV_RS06660 tRNA (adenosine(37)-N6)-dimethylallyltransferase MiaA
BOV_RS00995 thiaminase II
BOV_RS04130 phosphoribosylformylglycinamidine synthase subunit PurL
BOV_RS04155 phosphoribosylaminoimidazolesuccinocarboxamide synthase
BOV_RS03500 phosphoribosylglycinamide formyltransferase
BOV_RS00690 bifunctional uridylyltransferase/uridylyl-removing protein
BOV_RS02130 phosphoribosylamine--glycine ligase
BOV_RS02280 amidophosphoribosyltransferase
BOV_RS03505 phosphoribosylformylglycinamidine cyclo-ligase
BOV_RS08585 bifunctional phosphoribosylaminoimidazolecarboxamide formyltransferase/inosine monophosphate cyclohydrolase
BOV_RS07900 hypothetical protein
BOV_RS08630 lytic murein transglycosylase
BOV_RS09045 tRNA (guanosine(37)-N1)-methyltransferase TrmD
BOV_RS06065 alpha/beta hydrolase
BOV_RS08265 5-(carboxyamino)imidazole ribonucleotide mutase
BOV_RS09735 tRNA uridine-5-carboxymethylaminomethyl(34) synthesis enzyme MnmG
BOV_RS09740 tRNA uridine-5-carboxymethylaminomethyl(34) synthesis GTPase MnmE
BOV_RS10110 phosphate ABC transporterpermease protein PstA
BOV_RS04795 RluA family pseudouridine synthase
BOV_RS01785 saccharopine dehydrogenase
BOV_RS06060 serine O-acetyltransferase
BOV_RS04630 ATP-dependent helicase
BOV_RS05445 L D-transpeptidase
BOV_RS03110 AMP nucleosidase
BOV_RS09535 heavy metal translocating P-type ATPase
BOV_RS09615 D-alanyl-D-alanine carboxypeptidase
BOV_RS15320 elongation factor 4
BOV_RS04930 DUF922 domain-containing protein
BOV_RS03230 GTP pyrophosphokinase rsh
BOV_RS05340 HAMP domain-containing histidine kinase
BOV_RS06975 alpha/beta hydrolase
BOV_RS14345 type I restriction endonuclease subunit R
BOV_RS02060 tRNA1(Val) (adenine(37)-N6)-methyltransferase
BOV_RS04850 glutathione-disulfide reductase
BOV_RS01700 large conductance mechanosensitive channel protein MscL
BOV_RS00245 L D-transpeptidase
BOV_RS06915 N-acetylmuramoyl-L-alanine amidase
BOV_RS04900 MCE family protein
BOV_RS10170 PhoH family protein
BOV_RS00145 putative sulfate exporter family transporter
BOV_RS00475 dihydroxy-acid dehydratase
BOV_RS01830 (S)-ureidoglycine aminohydrolase
BOV_RS05655 MerR family DNA-binding transcriptional regulator
BOV_RS06655 acetolactate synthase 3 large subunit
BOV_RS05540 RIP metalloprotease RseP
BOV_RS06685 protease modulator HflC
BOV_RS10675 hypothetical protein
BOV_RS00460 heme exporter protein C
BOV_RS13135 cytochrome c biogenesis protein CcdA
BOV_RS02505 hypothetical protein
BOV_RS07830 disulfide bond formation protein B
BOV_RS05580 DUF1849 domain-containing protein
BOV_RS03700 RNA degradosome polyphosphate kinase
BOV_RS03540 L-lactate permease
BOV_RS06675 PDZ domain-containing protein
BOV_RS09555 cobaltochelatase subunit CobS
BOV_RS10595 amino acid permease
BOV_RS07420 DUF58 domain-containing protein
BOV_RS02300 class I SAM-dependent methyltransferase
BOV_RS07425 DUF4159 domain-containing protein
BOV_RS07430 hypothetical protein
BOV_RS15415 cardiolipin synthetase
BOV_RS08905 hypothetical protein
BOV_RS08900 heme biosynthesis protein HemY
BOV_RS14115 acyl-CoA dehydrogenase
BOV_RS09550 cobaltochelatase subunit CobT
BOV_RS13960 glutamine synthetase
BOV_RS09365 TIGR02302 family protein
BOV_RS09775 maf-like protein
BOV_RS11600 ribulokinase
BOV_RS12590 thiol reductant ABC exporter subunit CydD
BOV_RS10720 oligoendopeptidase F
BOV_RS04905 hypothetical protein
BOV_RS15345 NAD(P)-dependent oxidoreductase
BOV_RS07415 MoxR family ATPase
BOV_RS08595 NAD-glutamate dehydrogenase
BOV_RS14750 histidine utilization repressor
BOV_RS08845 phosphoenolpyruvate--protein phosphotransferase
BOV_RS08345 hypothetical protein
BOV_RS07285 peptidoglycan editing factor PgeF
BOV_RS10290 peptidase P60
BOV_RS00895 methionine synthase
BOV_RS04615 glycosyltransferase family 2 protein
BOV_RS01715 glycosyl transferase
BOV_RS02700 mannose-1-phosphate guanylyltransferase/mannose-6-phosphate isomerase
BOV_RS07170 glycosyltransferase family 1 protein
phosphomannomutase
BOV_RS07180 DUF1176 domain-containing protein
BOV_RS10190 XRE family transcriptional regulator
BOV_RS01370 DUF2948 domain-containing protein
BOV_RS05330 Trk system potassium transporter TrkA
BOV_RS03070 hypothetical protein
BOV_RS10970 ChbG/HpnK family deacetylase
BOV_RS10975 glycosyltransferase family 39 protein
BOV_RS14735 D-amino acid dehydrogenase small subunit
BOV_RS01585 two-component sensor histidine kinase
BOV_RS01125 sugar ABC transporter substrate-binding protein
BOV_RS02550 MBL fold metallo-hydrolase
BOV_RS13775 glycine dehydrogenase
BOV_RS13785 glycine cleavage system aminomethyltransferase T
BOV_RS06590 PhzF family phenazine biosynthesis protein
BOV_RS10125 phosphate regulon transcriptional regulatory protein PhoB
BOV_RS07865 RluA family pseudouridine synthase
BOV_RS12140 FadR family transcriptional regulator
BOV_RS14100 helix-turn-helix transcriptional regulator
BOV_RS03705 exopolyphosphatase
BOV_RS10070 SMP-30/gluconolactonase/LRE family protein
BOV_RS09630 NAD(P)-dependent oxidoreductase
BOV_RS11585 sugar ABC transporter ATP-binding protein
BOV_RS11580 ABC transporter substrate-binding protein
BOV_RS11590 sugar ABC transporter permease
BOV_RS11815 class 1b ribonucleoside-diphosphate reductase subunit alpha
BOV_RS01680 prolyl aminopeptidase
BOV_RS10060 orotidine-5'-phosphate decarboxylase
BOV_RS12310 glycerol kinase
BOV_RS13210 aspartate carbamoyltransferase
BOV_RS13215 dihydroorotase
Increasing Fitness Score
Decreasing Fitness Score
3.4
3.0
2.6
2.2
1.8
1.4
1.0
0.6
0.2
-0.2
-0.6
-1.0
-1.4
-1.8
-2.2
-2.6
-3.0
-3.4

**Supplemental Figure 1. Fitness profile of *B. ovis* transposon insertion mutants as a function of growth phase.**

Heat map of fitness scores of 118 genes with t-like significance score  |4|. Genes were grouped by hierarchical clustering. Each column is a point during the growth curve ($OD_{600}$, indicated at the top of the heatmap). Each row is a gene. Gene locus tags and annotated functions are indicated on the left.

**A**

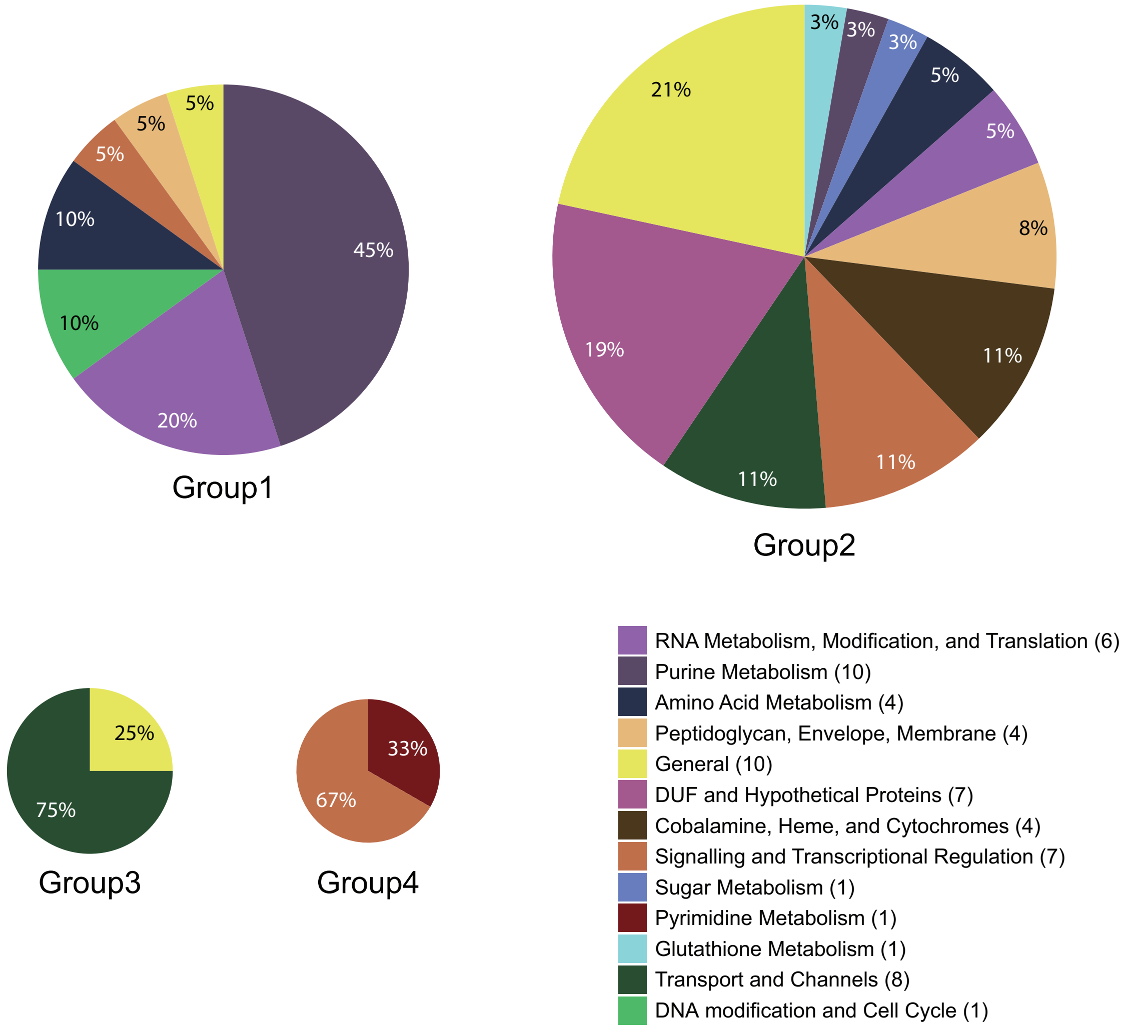


**B**

| Annotated product | Locus ID | Average fitness score | | | | Group |
|---|---|---|---|---|---|---|
| | | 0.05 | 0.12 | 0.9 | 2.4 | |
| serine O-acetyltransferase | *BOV_06060* | - 0.18 | - 0.11 | - 1.65 | - 3.31 | 1 |
| AMP-nucleosidase | *BOV_03110* | - 0.79 | - 1.16 | - 0.99 | - 3.06 | 1 |
| bifunctional (p)ppGpp synthetase/guanosine-3',5'-bis(diphosphate) 3'-pyrophosphohydrolase | *BOV_03230* | - 1.52 | - 0.18 | - 1.78 | - 2.65 | 2 |
| saccharopine dehydrogenase family protein | *BOV_01785* | - 0.2 | - 0.32 | - 1.22 | - 2.45 | 1 |
| MerR family DNA-binding transcriptional regulator | *BOV_05655* | - 1.33 | - 1.29 | - 1.72 | - 2.43 | 2 |
| alpha-beta hydrolase | *BOV_06975* | - 0.52 | - 0.93 | - 1.73 | - 2.21 | 2 |

**Supplemental Figure 2. Functional classification of *B. ovis* genes whose disruption significantly impacts fitness during growth in Brucella broth.**
A**)** Gene function pie chart of the four clusters of genes identified in **Fig. 1,** grouped based on predicted function. Area of each pie chart is proportional to the number of genes in that group. DUF = Domain of Unknown Function. **B)** Subset of genes from **Fig. 1A** with fitness score values □ |2| in stationary phase.

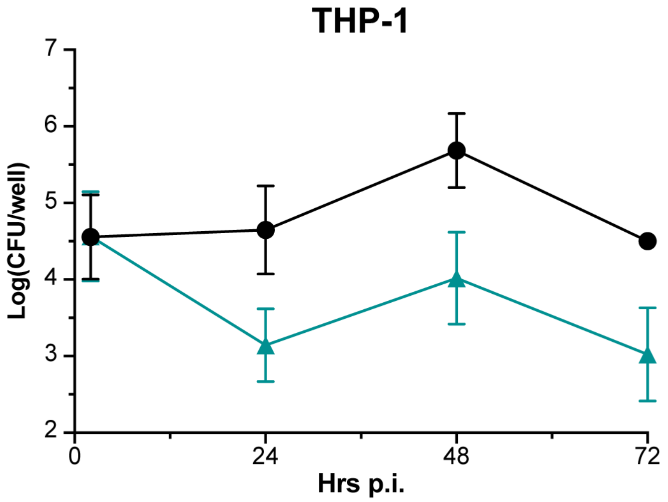


**Supplemental Figure 3. Recovered CFUs decrease at 72 hrs post infection**
Colony forming units of wild type *B. ovis* (WT, black circles) and Δ*cysE* (teal triangles) cells recovered from infected THP-1 cells at 2, 24, 48, and 72 hours post infection (Hrs p.i.). Error bars represent SEM for three independent experiments.

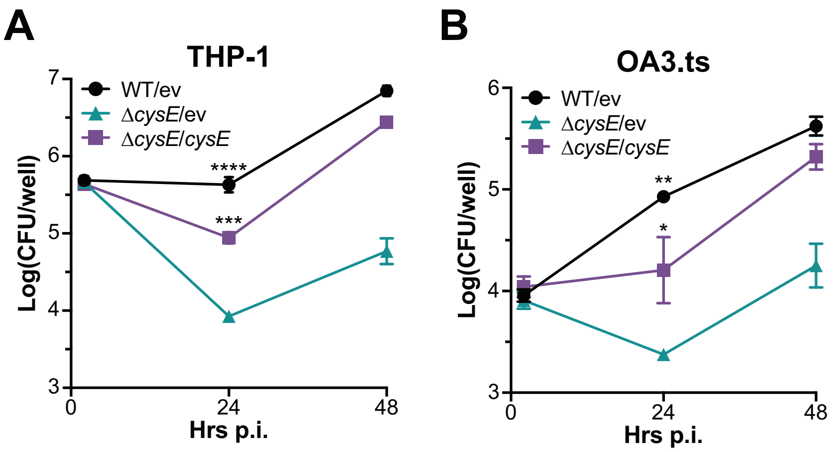


**Supplemental Figure 4. Genetic complementation of THP-1 and OA3.ts infection**
$Log_{10}$ colony forming units (CFU) per well of WT/pSRK-EV (WT/ev, black circles), Δ*cysE*/pSRK-EV (Δ*cysE*/ev teal triangles), and Δ*cysE*/pSRK-cysE(Δ*cysE*/*cysE*, purple diamonds) isolated from infected THP-1 (**A**) or OA3.ts mammalian cells (**B**). Significance at the 24 hr time point was assessed using one-way ANOVA (Tukey's multiple comparison test). Asterisks indicate p-values compared to Δ*cysE* or Δ*cysE*/ev: * $p < 0.05$; ** $p < 0.01$; *** $p < 0.001$, **** $p < 0.0001$. Infections were repeated 3 -5 independent times. Error bars represent standard error within the representative experiment.

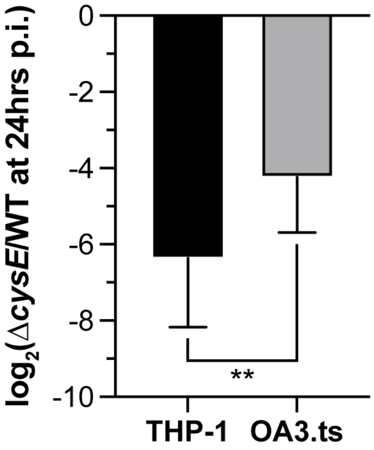


**Supplemental Figure 5. *B. ovis* Δ*cysE* is more attenuated in THP-1 human macrophage-like cells than in ovine testis OA3.ts cells.**

Ratio of recovered CFUs of *B. ovis* Δ*cysE* relative to WT at 24 hrs post infection (p.i.) in macrophage-like human THP-1 cells and in the sheep testis epithelial cell line, OA3.ts. p-value was calculated using a two-tailed unpaired t-test, ** $p<0.01$. Error bars represent SD of four independent experiments in each cell line.

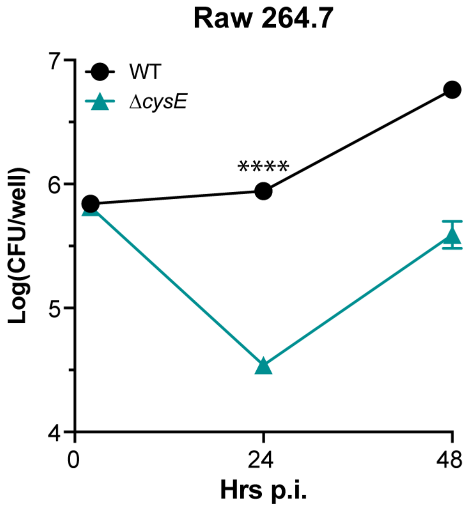


**Supplemental Figure 6. *B. ovis* Δ*cysE* strain is attenuated in Raw 264.7 macrophages**

Colony forming units of wild type (black circles) and Δ*cysE* (teal triangles) *B. ovis* cells recovered after infection of Raw 264.7 murine macrophages. p-value calculated using an unpaired t-test for the 24 hr time point. Error bars represent SEM for four independent repetitions of the experiment.
